# A large kinome in a large cell: *Stentor coeruleus* possesses highly expanded kinase families and novel domain architectures

**DOI:** 10.1101/168187

**Authors:** Sarah B. Reiff, Wallace F. Marshall

## Abstract

**Background:** *Stentor coeruleus* is a large ciliated protist, about 1mm in length, with the extraordinary ability to fully regenerate each fragment after being cut into pieces, perfectly restoring cell polarity and morphology. Single-cell regeneration in *Stentor* remains one of the greatest long-standing mysteries of biology, but the recently published *Stentor* genome now enables studies on this organism at the molecular and genetic levels. Here we characterize the complete complement of kinases, or kinome, of *Stentor*, in order to begin to understand the signaling capacities that underlie *Stentor*’s unique biology.

**Results:** The genome of *S. coeruleus* contains over 2000 kinases, representing 6% of the predicted proteome. Classification of the kinase genes reveals large expansions in several kinase families, particularly in the CDPKs, the DYRKs, and in several mitotic kinase families including the PLKs, NEKs, and Auroras. The large expansion of the CDPK and DYRK kinase families is an unusual feature of the *Stentor* kinome compared to other ciliates with sequenced genomes. The DYRK family in *Stentor*, notably, contains only a single pseudokinase which may suggest an important role in *Stentor* growth and survival, while the smaller PEK family contains a novel pseudokinase subfamily. The *Stentor* kinome also has examples of new domain architectures that have not been previously observed in other organisms.

**Conclusion:** Our analysis provides the first gene-level view into the signaling capabilities of *Stentor* and will lay the foundation for unraveling how this organism can coordinate processes as complex as regeneration throughout a giant cell.

## Background

*Stentor coeruleus* stands out for its extremely large size and highly complex behavioral repertoire. The cell is 1mm long, with an oral apparatus at the anterior end that it uses for filter feeding, and precisely arranged ciliary rows that run from the anterior to the posterior (Figure 1). Despite being a unicellular organism, *Stentor* has the ability to fully regenerate after being cut in pieces in a way that perfectly preserves the original cell polarity and morphology [1]. *Stentor* can also respond to light and mechanical stimuli by changing swimming direction or by contracting. This contractile response is subject to habituation [2], even though this phenomenon is more often associated with multicellular nervous systems. The wide range of behaviors suggests a rich intracellular signaling capacity, especially since the cell is also so large, requiring signals to propagate over long intracellular distances.

**Figure 1.**
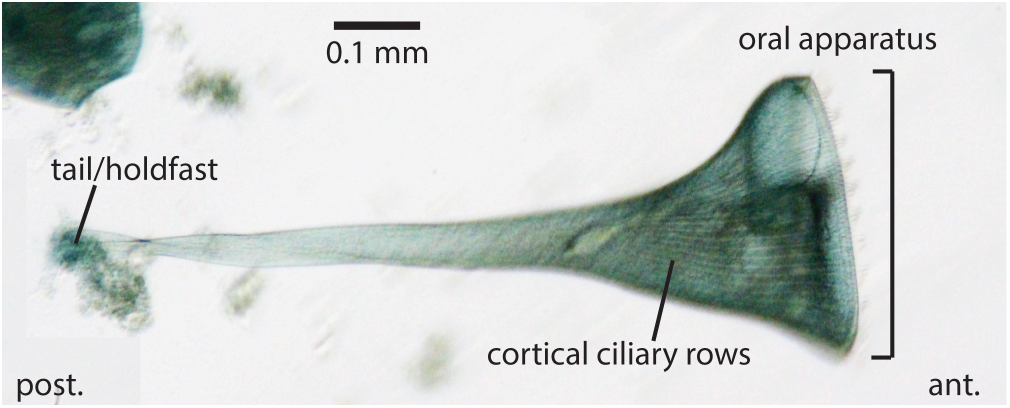
Stentor coeruleus cell structure. Micrograph of a single *Stentor* cell with a few cortical features labeled. Anterior (ant.) and posterior (post.) are indicated.

Recently our group has sequenced and assembled a draft genome for *S. coeruleus* [3], now publicly available (http://stentor.ciliate.org/). The 83 Mb genome has a high gene density, encoding over 34,000 genes. It is also quite intron-poor, and amazingly the vast majority of introns are only 15-16 bp in length, making *Stentor* the organism with the smallest introns known to date. The release of the genome presents new opportunities for studying the molecular details behind *Stentor* regeneration and behavior. How might the complexity of *Stentor* behavior, habituation, and regeneration be manifested in terms of the diversity of signaling proteins encoded by the genome?

Protein kinases are ubiquitous throughout biology and play key roles in all kinds of cell signaling pathways. Found in all domains of life, they have achieved a rich diversity through the course of evolution. Eukaryotic protein kinases (ePKs) share a common domain fold [4] and are classified into the following 7 main groups, each of which contain several related families and subfamilies: AGC, CAMK, CK1, CMGC, RGC, STE, TK, and TKL. ePKs that aren’t related to any of these groups are classified as “Other” kinases, and include some well-known kinases families like Aurora kinases and Polo-like kinases. In addition to ePKs, there are also atypical protein kinases, which are protein kinases that don’t share the typical kinase domain architecture characteristic of ePKs. These include the histidine kinases as well as the RIO, ABC, Alpha, and PIKK families. The evolutionary history of kinase families is closely coupled to the evolution of developmental and regulatory processes, such that the kinome of an organism is important for understanding the evolution of a cell’s behavior.

*Stentor* belongs to the ciliates, a protist phylum characterized by complex cortical ultrastructure and a diverse range of behaviors. While the kinases of metazoa, *Arabidopsis*, and many unicellular organisms tend to make up around 2-3% of protein coding genes [5–7], ciliates tend to have larger kinomes than most other organisms. The kinases of *Tetrahymena* and *Paramecium* make up 4% and 7% of the predicted proteomes, respectively [8, 9]. The expansion of kinase families in ciliates may suggest a need for multiple distinct but related kinase functions in regulating the behaviors of these large and complex free-living cells.

As a first step towards dissecting signaling in *Stentor*, we characterize the kinome of *S. coeruleus*, with the idea that most important cellular processes are likely to be regulated by kinases. We find that the *Stentor* genome encodes over 2000 kinases, with expansions in several different kinase families, reflecting the elaborate signaling needs of this cell.

## Materials and Methods

### Identification and classification of kinases and other proteins

Profile HMMs were downloaded for all the kinase groups, families, and subfamilies in Kinbase (http://kinase.com/web/current/kinbase/). These were compared to all the *Stentor* gene models (available at stentor.ciliate.org) using the hmmsearch tool in HMMER v3.1 (http://hmmer.org/), using the following command:

> *hmmsearch* -*E 0.001* –*cpu 16* –*noali* –*seed 544* –*tblout kinases_hmmsearch_tab.txt* -*o kinases_hmmsearch.txt kinasedomains.hmm Stentor_proteome.fasta*

Gene models with partial kinase domains were compared to the *S. coeruleus* genomic locus and vegetative RNA-seq data [3]. In 24 cases we were able to improve the gene models, resulting in full kinase domains, which have now been updated on the *Stentor* genome database (stentor.ciliate.org). All *S. coeruleus* kinase hits were then verified with BlastP [10] using a custom blast database of all kinase domains in Kinbase. If the highest scoring BlastP hit matched the highest scoring profile HMM, the gene was classified accordingly, at either the subfamily, family, or group level. For those that didn’t match, we used an all-against-all blastp search to cluster them into either pre-existing kinase families or new *Stentor*-specific kinase families, or as unique. Note that in KinBase, for the *Tetrahymena* genes that are classified into the Ciliate-E2b kinase subfamily, the majority of these genes were later reannotated as PKG kinases in the *Tetrahymena* genome database. Thus, *Stentor* genes that were matches for the Ciliate-E2b family have been classified as PKG family kinases here. Additionally, since the published kinome analysis of *Paramecium* [8] only classifies eukaryotic protein kinases at the group level, we have used this same approach to classify *P. tetraurelia* kinases into families and subfamilies as well as to identify atypical protein kinases (Additional File 1 Table S2), so that we could make direct comparisons.

To identify other types of proteins, namely cyclins, GPCRs, and guanylate cyclases, we used a similar HMMER/BlastP approach. For searches with hmmsearch, profile HMMs were downloaded from Pfam for the following Pfam domains: PF00134, PF02984, and PF08613 for cyclins; PF00211 for adenylate/guanylate cyclases; PF02116, PF02117, PF02175, PF10192, PF10324, PF10292, PF10316, PF10317, PF10318, PF10320, PF10321, PF10323, PF10326, PF10328, PF12430, PF08395, PF10319, PF10322, PF10325, PF10327, PF14778, PF00001, PF00003, PF00002, PF02076, PF03125, PF01036, PF05462, PF11710, PF02118, PF03383, PF03402 for GPCRs. The pfam gathering cutoff for each profile HMM was used as the reporting threshold.

### Identification of conserved tyrosine phosphorylation sites

For the MAPK family, CDC2 subfamily, and H2A family proteins in Stentor, sequences were aligned in MEGA7 using MUSCLE [11] with the UPGMB clustering method and default parameters. Sequences were then checked for the following conserved residues in the alignment: Y204 from human ERK1 in the MAPK alignment, Y15 from *S. pombe* CDC2 in the CDC2 alignment, and Y57 from human H2A in the H2A alignment.

### Analysis of domain architecture

Profile HMMs of all the domains in pfam-A were downloaded from ftp://ftp.ebi.ac.uk/pub/databases/Pfam/releases/Pfam27.0/. *Stentor* kinase genes were compared to these with the hmmsearch tool in HMMER, using the pfam gathering cutoff as the reporting threshold, with the following command:

> *hmmsearch* –*cut_ga* –*cpu 16* –*noali* –*seed 544* –*domtblout kinases_pfam_tab.txt* -*o kinases_pfam.txt Pfam-A.hmm Stentor_kinases.fasta*

### Phylogenetic analysis

To generate the ePK tree, we removed all the atypical kinases, leaving only the *Stentor* ePK sequences, and then removed genes with truncated kinase domains (less than 100 amino acids). These sequences were then initially aligned to the pfam PKinase domain profile hmm using the hmmalign tool in HMMER3. The resulting alignment was trimmed with TrimAl [12] and further aligned using SATe II [13]. The final alignment was then used in RAxML v8.2 [14] to generate a maximum likelihood (ML) tree under the WAG amino acid substitution model and gamma model of rate heterogeneity with 200 bootstrap replicates. Code for alignment, trimming, and ML analysis is available at https://github.com/sbreiff/StentorKinome. The resulting tree (Additional File 2) was visualized using FigTree (http://tree.bio.ed.ac.uk/software/figtree/).

The *Stentor* kinase alignment was also used to identify kinases with catalytic site substitions. We looked for substitutions in any of the following three sites from the Pkinase domain consensus: K30, D123, D141, which correspond to the lysine and aspartate residues in the conserved motifs VAIK, HRD, and DFG, respectively.

To generate the PEK tree, kinase domains from human, mouse, fly, yeast, Dictyostelium, and Tetrahymena sequences were downloaded from Kinbase, and *Stentor* and *Paramecium* sequences were aligned to these using MUSCLE in MEGA7. The resulting alignment was then trimmed with TrimAl using the “automated1” heuristic. The trimmed alignment was then analyzed in RAxML under the LG substitution model and gamma model of rate heterogeneity, with 250 bootstrap replicates, using the following command:

> *raxmlHPC*-*PTHREADS*-*AVX* –*T 16* –*f a* –*m PROTGAMMAAUTO* –*x 12345* –*p 12345* -# *autoMRE* –*s PEKalign_trim.fasta* –*n PEKtree*

The resulting tree was visualized in Figtree. The second pseudokinase domains of metazoan GCN2 were used as an outgroup to root the tree.

## Results

### The *Stentor coeruleus* genome contains over 2000 kinases

We used HMMER3 to search for kinase domains in the 34,506 *Stentor* protein-coding genes. In total we identified 2057 kinase genes (Additional File 1 Table S1), equaling 6.0% of protein-coding genes in *Stentor*. Although this is a much higher complement of kinases than most other eukaryotes, it is comparable with other ciliates, which tend to have higher numbers of kinases (Table 1). Using BlastP to confirm identities of hits, we also classified the *Stentor* kinases into kinase groups, families, and subfamilies. While most HMM and BlastP top hits corresponded, in roughly 10% of cases, the best HMM hit and best BlastP hit did not match, so we utilized blast-based clustering to classify these. Some of these could be classified into pre-existing families, but 66 were unique, and 134 were grouped into provisional *Stentor*-specific families. 10 contained partial kinase domains that were too small to classify. Maximum likelihood phylogenetic analysis recovers groupings based on our classifications (Figure 2).

**Figure 2.**
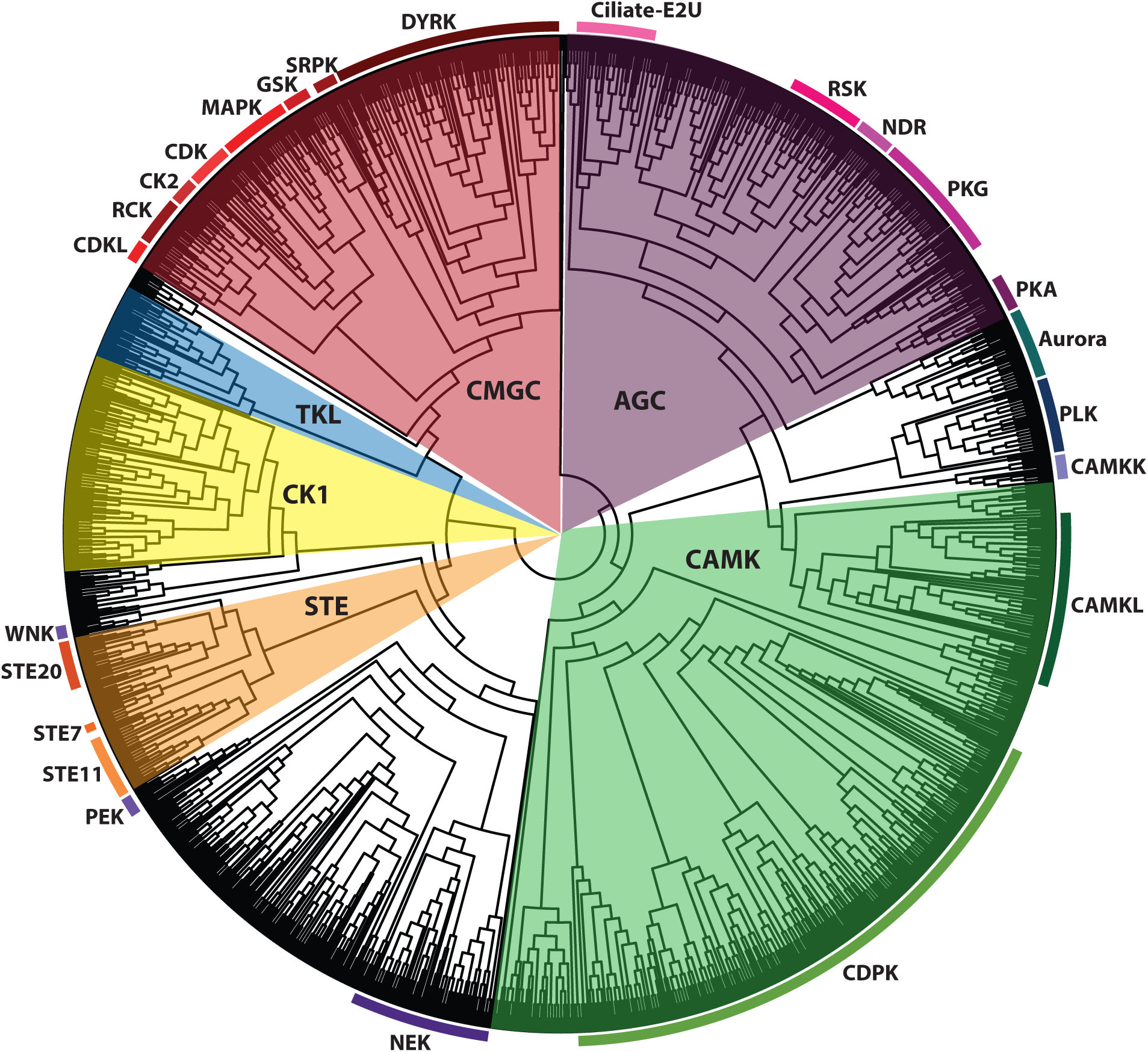
Dendrogram of Stentor ePKs. Shaded regions of the tree indicate kinase groups, and outer ring indicates selected kinase families.

**Table 1.** Summary table of kinome sizes.

| Organism | # of Kinases in Genome | Kinases as % of Protein-Coding Genes |
| --- | --- | --- |
| *Homo sapiens* | 518 | 2.7% |
| *Caenorhabditis elegans* | 438 | 2.1% |
| *Saccharomyces cerevisiae* | 113 | 2.0% |
| *Arabidopsis thaliana* | 1019 | 3.7% |
| *Dictyostelium discoideum* | 285 | 2.6% |
| *Phytophthora infestans* | 372 | 2.1% |
| *Giardia intestinalis* | 278 | 4.3% |
| *Plasmodium falciparum* | 89 | 1.6% |
| *Ichthyophthirius multifiliis*<sup>a</sup> | 671 | 8.3% |
| *Tetrahymena thermophila*<sup>a</sup> | 1068 | 4.0% |
| *Paramecium tetraurelia*<sup>a</sup> | 2841 | 7.2% |
| *Stentor coeruleus*<sup>a</sup> | 2057 | 6.0% |

<sup>a</sup> Ciliate.
Data sources: *H. sapiens* [5], *C. elegans* [15], *S. cerevisiae* [7], *A. thaliana* [6], *D. discoideum* [16], *P. infestans* [17], *G. intestinalis* [18], *P. falciparum* [19], *I. multifiliis* [20], *T. thermophila* [9], *P. tetraurelia* [8]. Note that the original *P. tetraurelia* analysis only examined eukaryotic protein kinases, but here we also include atypical protein kinases found in the genome.

As kinomes of the ciliates *T. thermophila* and *P. tetraurelia* have been previously characterized, some of the differences between these and *Stentor* will be highlighted (Table 2). When comparing kinase groups (Figure 3A), the biggest difference lies in the smaller proportion of “Other” kinases in the *Stentor* kinome, but this is largely due to the presence of fewer members of ciliate-specific families in *Stentor*. *Stentor* also has a smaller proportion of atypical kinases, but we find several kinase families that exhibited significant expansions in members in *Stentor* compared to other organisms (Figure 3B; Table 2). Some of the major kinase families in each group are discussed below.

**Figure 3.**
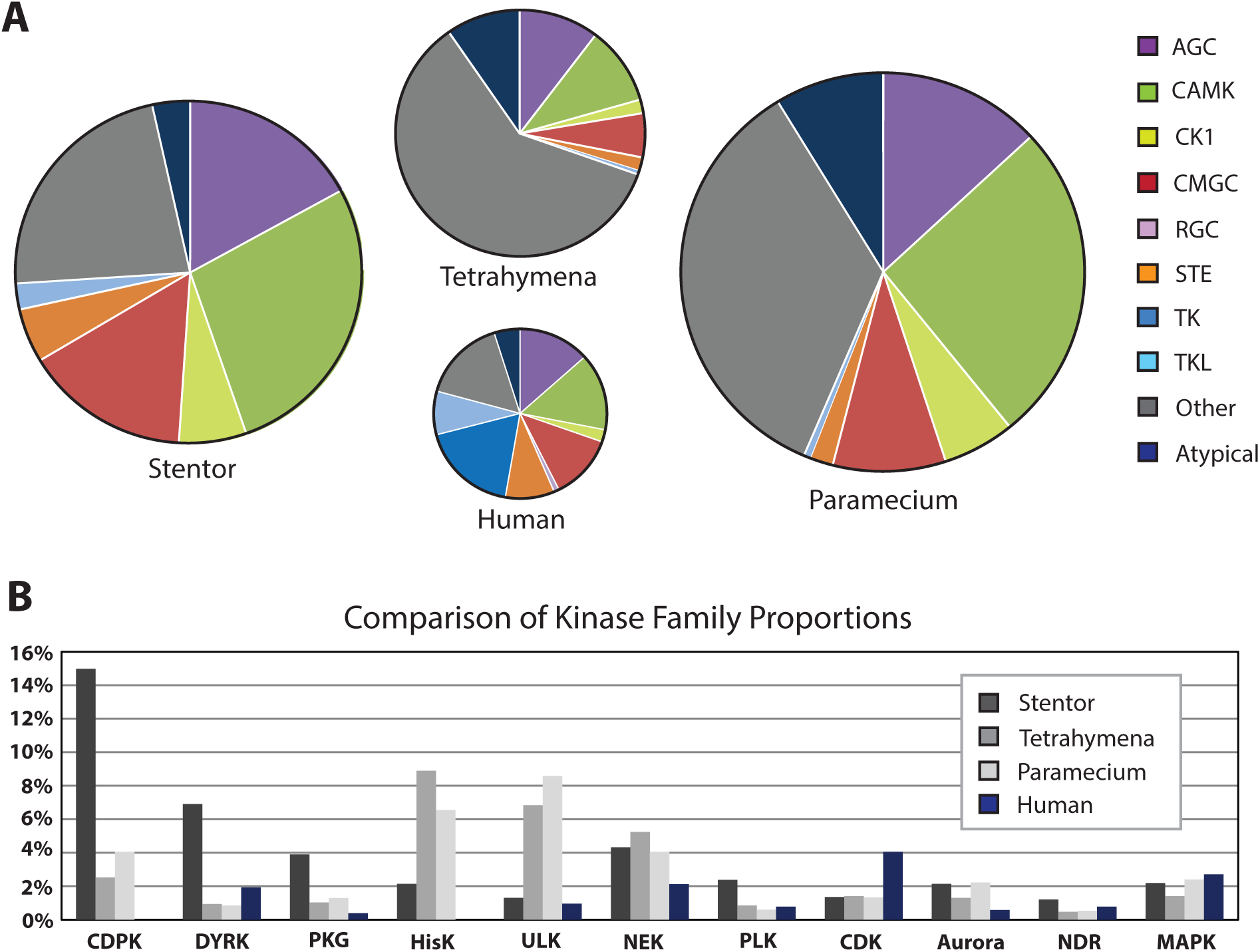
Comparison of kinomes across 4 species. **A**. Kinase groups shown as proportions of the total kinome in *S. coeruleus*, *T. thermophila*, *P. tetraurelia*, and *H. sapiens*. Size of circle is proportional to kinome size in each species. **B**. Bar graph comparison of 11 selected families in terms of proportion of kinome. Absolute numbers can be found in Table 2.

**Table 2.** Selected kinase families in Stentor compared to other model organisms.

| Group | Family | *Stentor* | *Tetrahymena* | *Paramecium* | Yeast | Human |
| --- | --- | --- | --- | --- | --- | --- |
| AGC | PKA | 23 | 4 | 23 | 3 | 5 |
|  | PKC | 0 | 0 | 0 | 1 | 9 |
|  | PKG | 79 | 7 | 37 | 0 | 2 |
|  | NDR | 25 | 5 | 15 | 4 | 4 |
|  | MAST | 0 | 3 | 19 | 0 | 5 |
|  | RSK | 48 | 5 | 18 | 3 | 8 |
| CAMK | CDPK | 308 | 27 | 115 | 0 | 0 |
|  | CAMKL | 107 | 22 | 96 | 12 | 20 |
| CMGC | DYRK | 142 | 10 | 24 | 1 | 10 |
|  | CDK | 28 | 15 | 38 | 7 | 21 |
|  | CDKL | 14 | 4 | 11 | 0 | 5 |
|  | CK2 | 17 | 3 | 16 | 2 | 2 |
|  | GSK | 18 | 7 | 28 | 4 | 2 |
|  | RCK | 31 | 8 | 23 | 1 | 3 |
|  | SRPK | 15 | 3 | 16 | 1 | 3 |
|  | MAPK | 45 | 15 | 68 | 6 | 14 |
| CK1 | CK1 | 130 | 20 | 164 | 4 | 7 |
| STE | STE7 | 5 | 4 | 9 | 4 | 7 |
|  | STE11 | 39 | 8 | 25 | 5 | 8 |
|  | STE20 | 29 | 7 | 10 | 7 | 30 |
| TKL |  | 46 | 8 | 15 | 0 | 43 |
| Other | Aurora | 44 | 15 | 63 | 1 | 3 |
|  | NEK | 89 | 56 | 115 | 1 | 11 |
|  | PLK | 49 | 9 | 17 | 1 | 4 |
|  | ULK | 27 | 73 | 244 | 1 | 5 |
|  | CDC7 | 0 | 1 | 7 | 1 | 1 |
|  | CAMKK | 16 | 9 | 18 | 4 | 2 |
|  | Wee | 2 | 2 | 4 | 1 | 3 |
|  | WNK | 10 | 2 | 8 | 0 | 4 |
|  | PEK | 13 | 4 | 9 | 1 | 4 |
| Atypical | HisK | 44 | 95 | 186 | 1 | 0 |
|  | Alpha | 3 | 9 | 20 | 0 | 6 |
|  | PIKK | 19 | 15 | 34 | 5 | 6 |
|  | ABC1 | 3 | 4 | 6 | 3 | 5 |
|  | RIO | 3 | 2 | 2 | 2 | 3 |
| All |  | 2057 | 1068 | 2841 | 113 | 518 |

Data sources: *Stentor* and *Paramecium* - this study; *Tetrahymena*, Yeast, Human – Kinbase.

### Atypical Kinases

*Stentor* encodes 72 genes for Atypical kinases. Because these possess a domain fold significantly different from the ePK domain, these could not be included in our phylogenetic analysis with the other kinases. Among the atypical kinases in *Stentor* are 3 ABC1 kinases, 3 Alpha kinases, 3 RIO kinases, 19 PIKKs and 44 Histidine kinases (HisKs). Many of these families are poorly understood, but HisKs are known to mainly function in two-component signaling systems and often serve to detect extracellular signals. When the *Tetrahymena thermophila* macronuclear genome was published, one of the notable findings in terms of the kinases encoded was a large expansion of histidine kinases [9]; the *Paramecium* kinome exhibits a similar pattern (Additional File 1 Table S2). While the number of HisKs in Stentor is higher than in most other eukaryotes, it is reduced compared to *Tetrahymena* and *Paramecium* (Table 2 and Figure 3B), even though the *Tetrahymena* kinome is smaller.

### AGC Group

Among the ePKs, *Stentor* contains a total of 348 AGC group kinases. This group was named for Protein Kinases A, G, and C, but also includes the NDR, RSK, and MAST families. Consistent with other ciliates, *Stentor* encodes no homologs of Protein Kinase C. However, there are 23 PKAs and 79 PKGs. The presence of multiple PKA and PKG orthologs suggests that *Stentor* uses cAMP and cGMP as upstream regulators of kinase activity. We therefore looked for other members of cAMP and cGMP signaling pathways in *Stentor* as well.

*Stentor* has 67 adenylyl/guanylyl cyclases (Additional File 1 Table S3). Cyclase domains are very similar between adenylyl and guanylyl cyclases so we were unable to distinguish between these two types of proteins, but we did find two main domain architectures among these 66. 26 of them consisted of a single guanylate cyclase (GC) domain, but 38 possessed a domain architecture consisting of a P-type ATPase domain followed by two GC domains. Both domain architectures are common in alveolates, the superphylum containing ciliates, but the latter is alveolate-specific [21]. In *Paramecium*, some of these guanylyl cyclases are activated in response to calcium signaling [22, 23].

In metazoa as well as other eukaryotes, cAMP signaling is often driven by G-protein coupled receptors (GPCRs), so we searched the *Stentor* gene models for these domains as well. We found that *Stentor* encodes 80 GPCRs. 66 of these appear similar to the cAMP receptor family of GPCRs originally described in *Dictyostelium*; the rest group with the secretin family, the rhodopsin family, another rhodopsin-like family, the abscisic acid family, a glucose receptor family, and the ODR4-like family of GPCRs (Additional File 1 Table S3). All of these GPCR families are present in other ciliates except for the ODR4-like family, from which *Stentor* contains two members.

With 67 cyclases, 80 GPCRs, and 102 combined PKAs and PKGs, it appears that cAMP and cGMP signaling pathways are expanded in toto, including both the kinases themselves and their upstream regulators, suggesting a capacity for the cell to transduce signals from a wide variety of inputs. Interestingly, we could not find any homologs of metazoan G proteins. This is consistent with other ciliates, however, and since other parts of the signaling pathway are present, it is possible that *Stentor* and other ciliates encode a G protein that is too divergent from metazoan G proteins to be found by homology-based approaches.

The NDR kinase family comprises 25 members in *Stentor*. Our group has previously shown that the kinase regulator Mob1 is a polarity marker and essential for proper growth and regeneration in *Stentor* [24]. In metazoa, Mob1 functions in the Hippo pathway that helps regulate cell proliferation and apoptosis. In this pathway, Mob1 interacts with LATS1/2, which are NDR family kinases [25]. In *Stentor* these may serve roles in establishment and maintenance of polarity during growth and regeneration, as have been shown for Mob1. The MAST family is another AGC family that bears sequence similarities to NDR kinases [26], and while MAST kinases are present in metazoa and other ciliates, we do not find any members in *Stentor*.

### CAMK Group

The CAMK group is mainly composed of calcium- and calmodulin-dependent kinases and totals 561 kinases in *Stentor*, or 27% of the kinome. Calcium signaling is known to be crucial to cell physiology, particularly in ciliates. In *Vorticella* its role in the rapid contraction of the stalk has been well-studied [27, 28] and in *Paramecium* along with contraction it plays a major role in a variety of processes, especially motility and exocytosis [29–31]. In *Stentor*, calcium flux is involved in oral regeneration as well as contraction and the photophobic response [32–34]. Several types of kinases can respond to calcium, including the CDPK (calcium dependent protein kinases), calmodulin dependent protein kinase (CaMK), and protein kinase C (PKC). *Stentor* possesses five kinases in the CaMK1 family, a family conserved in metazoa. The CDPK family, however, appears to have the most dramatic expansion of any kinase family in *Stentor*, with 308 members (Figure 3B). The CDPKs constitute a family of calcium-dependent but calmodulin-independent protein kinases that possess EF hands that bind calcium in a calmodulin-like fashion. They were originally discovered in plants [35], where they are mainly involved in stress and immune signaling. CDPKs are absent from metazoa and fungi, but are present in apicomplexan parasites, a sister group to the ciliates, where they help regulate a variety of processes including motility, host cell invasion, and life cycle transitions [36]. Most of the organisms that possess CDPKs including plants and other alveolates seem to lack PKC, which plays roles in calcium signaling in metazoa and yeast. *Stentor* does, however, encode homologs of proteins that function upstream of PKC in humans, such as phospholipase C and phosphoinositol-3 kinase, so perhaps CDPKs are filling this role in *Stentor* and other protists.

The CaMK-like (CAMKL) family kinases, which are related to other CAMKs but are not themselves calcium-dependent, total 107 members in *Stentor*. 42 of these belong to the 5’-AMP-activated kinase (AMPK) subfamily, which are known to be involved in energy homeostasis [37]. An additional 28 belong to the microtubule affinity-regulating kinase (MARK) subfamily. In metazoa, MARKs help regulate microtubule dynamics [38], so this expanded family may help modulate the extensive microtubule-based cortical patterning of *Stentor*.

### CMGC Group

The CMGC group is named for the CDK, MAPK, GSK, and CDKL families, and contains 317 kinases in *Stentor*, representing a higher proportion of the kinome than observed in other ciliates or in humans (Figure 3A).The CDK family contains the canonical cell cycle regulators of other systems. *Stentor* encodes 28 CDKs, including 11 members of the cdc2 family. Generally, CDKs are regulated by binding to different cyclins at different stages of the cell cycle. To determine whether this type of regulation is also conserved in *Stentor*, we used HMMER to identify genes with cyclin domains among the predicted proteome, and confirmed hits with BlastP. We find a total of 106 cyclin genes, including 25 that contain both the N-terminal and C-terminal cyclin domains (Additional File 1 Table S3). In addition, two of the CDK genes possess cyclin domains, as has been described in *Phytophthora infestans* [17]. Metazoan systems typically possess less than 30 different cyclins, so 106 represents a massive expansion in *Stentor*.

*Stentor* also contains a startling 142 members of the dual specificity YAK-related kinase (DYRK) family, which has the ability to phosphorylate tyrosine as well as serine and threonine. This comes to 6.9% of the kinome, compared to 10 members in *Tetrahymena* and 24 in *Paramecium* (0.9% of their respective kinomes) and 10 in humans (2.0%). In humans, Dyrk1A is thought to contribute to Down Syndrome, and Dyrk2 is involved in hedgehog signaling and cell cycle control [39–41]. Other DYRKs include PRP4 which is involved in mRNA splicing and Yak which negatively regulates the cell cycle in budding yeast and modulates G-protein signaling during growth in *Dictyostelium* [42–44]. Of the *Stentor* DYRK family members, 56 belong to the Dyrk2 subfamily, while there are 8 in the plant-like DyrkP subfamily, 5 in the Yak subfamily, and 1 PRP4 homolog, suggesting that multiple DYRK clades contributed to the DYRK family expansion in *Stentor*. The remaining 72 weren’t classified into subfamilies, but appear most similar to DYRK2.

Also among the CMGC group in *Stentor* are 15 SRPKs, 17 CK2s, and 31 RCKs. SRPKs typically phosphorylate SR-rich proteins and play important roles in regulating mRNA maturation [45, 46]. CK2s are implicated in a wide array of cellular functions, having over 300 targets in humans [47, 48]. RCKs generally aren’t well characterized, but the MOK and MAK subfamilies are implicated in the regulation of ciliogenesis [49]. In particular, null mutants of the *Chlamydomonas reinhardtii* MOK kinase LF4 result in abnormally long flagella [50].

Finally, there are also 18 GSKs and 14 CDKLs, as well as 45 MAPKs. MAP kinase signaling cascades are found throughout eukaryotes and play essential functions in signal transduction, typically linking an extracellular signal to an alteration in gene expression. The MAPK family in *Stentor* includes 13 members of the ERK subfamily and 8 members of the ERK7 subfamily.

### STE Group

The STE group is named for the yeast sterile kinases. *Stentor* contains 104 STE kinases, which represents a greater proportion of the kinome than in other ciliates but much lower than metazoa or *Dictyostelium* (Figure 3A) [5, 16]. In *Stentor* 5 STE kinases are in the STE7 family, 39 are in the STE11 family, and 29 are in the STE20 family. In other systems, these families contain kinases involved in the MAPK cascade, namely MAPK kinases (MEKs), MEK kinases (MEKKs), and MEKK kinases (MAP4Ks). The MEKs are subfamilies of the STE7 kinase family and the MEKKs are subfamilies of the STE11 kinase family, while the MAP4Ks are members of the KHS subfamily of the STE20 kinases. *Stentor* kinases in the STE7 and STE11 families appear to have homology to MEKs and MEKKs by BLAST, and may serve these roles. However, although *Stentor* also contains STE20 family kinases, these do not seem to bear great similarity to MAP4Ks specifically.

Of the *Stentor* STE20 kinases, 2 belong to the MST subfamily. These are orthologs of the Drosophila Hippo kinase, which serves to activate NDR kinases in the Mob1 pathway mentioned earlier. In *Tetrahymena*, an MST homolog has been shown to be important for proper placement of the cell division plane [51].

### CK1 and TKL Groups

Consistent with other ciliates, *Stentor* does not encode any kinases from the RGC or TK group, which consist of receptor guanylyl cyclases and tyrosine kinases, respectively. Indeed, to this point these groups have mainly been observed in metazoa and choanoflagellates. *Stentor* does, however, contain 46 TKL (TK-like) kinases, which have a similar domain fold to the TKs but phosphorylate on serine/threonine. While this group is larger in *Stentor* than other ciliates, it represents a much smaller proportion of the kinome than in metazoa or *Dictyostelium* (Figure 2A) [5, 16]. There are also 130 members of the CK1 group, which consists only of the casein kinase 1 family in *Stentor*.

### “Other” Kinases

These comprise kinase families that aren’t related to any of the previous groups, and total 460 kinases in *Stentor*. This includes some major mitotic kinase families, such as Aurora kinases, PLKs, and NEKs. The aurora kinases are well studied for their roles in progression through mitosis, and *Stentor* encodes 44 aurora kinase genes. In addition, *Stentor* also contains 49 PLKs and 89 NEKs. These families are known to play roles in both the cell cycle and in ciliogenesis in metazoa. With a highly patterned cell cortex containing tens of thousands of cilia, expansions in these families may allow *Stentor* to keep up with the ciliogenesis needs of the cell throughout the cell cycle. Along with the previously mentioned NDRs and CDKs, mitotic kinases in *Stentor* often seem to exhibit expanded family sizes (Figure 3B).

A notable finding in the *Tetrahymena* kinome was a large expansion of the ULK kinases [9], which is also true of *Paramecium* (Figure 3B, Table 2). The functions of the ULK kinases are not very well understood, though in other systems they are reported to be involved in a variety of activities including hedgehog signaling, motile ciliogenesis, autophagy, and ER-to-Golgi trafficking [52–54]. In contrast to the other ciliates mentioned, *Stentor* only contains 27 ULKs, less than half the number found in *Tetrahymena*.

The kinase analysis in the *Tetrahymena* genome delineated several putative ciliate-specific kinase families [9]. Many of these families appear not to be conserved in *Stentor*, but we do find 7 kinases in the Ciliate-A9 family, 11 in the Ciliate-C1 family, 3 in the Ciliate-C4 family, and 52 in the Ciliate-E2-Unclassified subfamily (Additional File 1 Table S1). The “Other” kinases in *Stentor* also include 10 WNK kinases, 16 CAMKKs, 13 PEKs, and 2 Wee kinases.

## Tyrosine Phosphorylation and Dual Specificity Kinases

While *Stentor* lacks canonical TKs, other organisms lacking TKs do contain phosphotyrosine, usually produced by cryptic tyrosine kinases or dual-specificity kinases from other kinase groups. Above we described a large expansion in DYRKs in *Stentor*. However, while previously studied DYRKs autophosphorylate on tyrosine residues, they only transphosphorylate on serine or threonine residues [55]. Thus, this family is not expected to drive tyrosine phosphorylation of other substrates in *Stentor*, even though they are highly expanded.

Other kinase families have been shown to exhibit dual specificity in their transphosphorylation activities, including STE7, Wee, and CK2. In yeast and metazoa, Wee1 phosphorylates the tyrosine residue of the motif GxGTYG on CDK1 (CDC2), leading to inactivation [56, 57]. *Stentor* has 11 kinases in the CDC2 subfamily, so we performed an alignment and looked for this particular motif. This motif was conserved in 10 of the CDC2 kinases (Figure 4A), suggesting that the substrate for the *Stentor* Wee kinases is conserved, and likely its tyrosine phosphorylation activity as well. The only CDC2 kinase that doesn’t conserve this sequence is divergent across the whole kinase domain including at positions of critical catalytic sites, and happens to be one of the CDKs that possess two cyclin domains at the C-terminus (Figure 4B), raising the possibility that it may act primarily as a cyclin that uses a pseudokinase domain as a scaffold.

**Figure 4.**
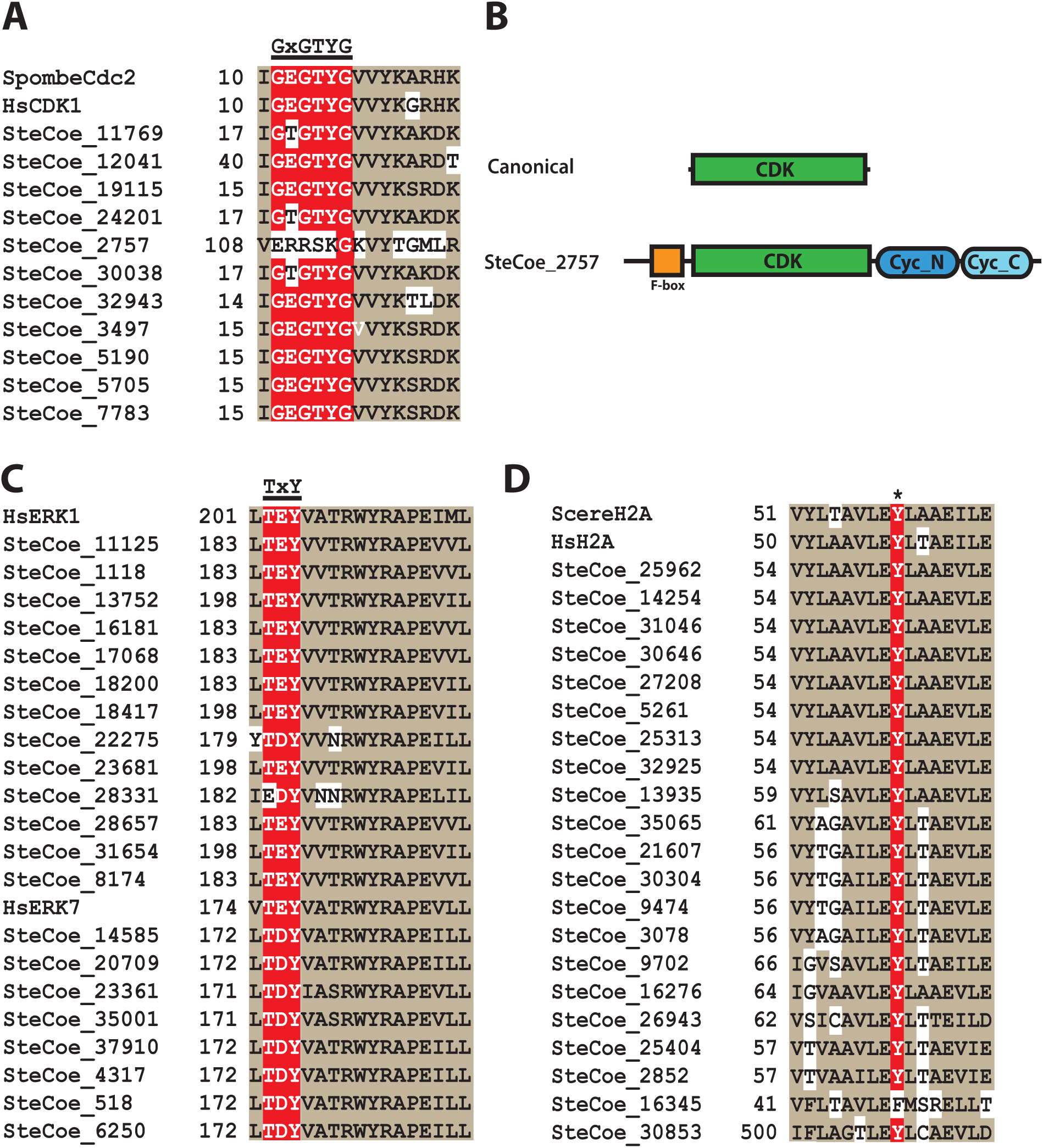
Conserved tyrosine phosphorylation sites in *Stentor* proteins. **A:** Alignment of *S. pombe* cdc2 and *H. sapiens* CDK1 with *Stentor* homologs at the GxGTYG tyrosine phosphorylation motif. **B:** Comparison of canonical cdc2 and SteCoe_2757 domain architectures. Drawn to scale. **C:** Alignment of *H. sapiens* ERK1 and ERK7 sequences with *Stentor* homologs at the TxY phosphorylation motif. **D:** Alignment of *S. cerevisiae* and *H. sapiens* H2A sequences with *Stentor* homologs at the Y57 phosphorylation site.

The STE7 family, also known for dual specificity transphosphorylation, consists of MEKs (MAP kinase kinases). It is known in metazoa that MAPKs are activated when the threonine and tyrosine residues in a TxY motif in the activation loop are phosphorylated [58, 59]. In ERK1/2, this sequence is typically TEY, but in other MAPK subfamilies the middle residue can be different [58, 59]. We aligned all the *Stentor* MAPKs and looked for conservation of this motif. The TxY activation loop was found to be present in 31 of the 45 *Stentor* MAPKs, most commonly as TEY or TDY. This number includes 12 of 13 ERK subfamily kinases and all 8 ERK7 subfamily kinases (Figure 4C). Conservation of a major STE7 kinase substrate suggests conserved tyrosine phosphorylation ability.

While the function of Wee kinases and MEKs have been well studied, CK2 kinases are much more pleiotropic and not as well understood. They are ubiquitous, constitutively expressed, and thought to act on hundreds of substrates, many of which seem to be involved in transcriptional activity and the spliceosome [48, 60]. For most substrates, CK2 probably phosphorylates on serine or threonine residues, but recently CK2 was found to phosphorylate histone H2A on Y57 to help regulate transcriptional elongation [61]. We searched the *Stentor* predicted proteome for homologs of H2A and identified 21 proteins with an H2A domain. We aligned these to the human H2A protein to look for conservation of Y57, and found that 20 of the 21 *Stentor* proteins had this residue conserved, suggesting that the tyrosine phosphorylation ability of CK2 is conserved in *Stentor* (Figure 4D). We thus infer from this analysis of kinase and substrate sequences that *Stentor* is likely to use tyrosine phosphorylation of substrates as a regulatory mechanism, relying on dual specificity kinases.

## Novel domain architectures in the *Stentor* kinome

Highly expanded kinase families create potential for diversification and subfunctionalization in its members. Gene fusions could potentially result in acquisition of additional protein domains that could facilitate greater functional diversity within a kinase family. We used HMMER3 to search the *Stentor* kinases for additional pfam domains. Many of the protein domains uncovered in the kinases helped confirm classifications: the polo-like kinase homologs had polo domains, for example, while PKGs had nucleotide-binding domains and CDPKs possessed EF hands for calcium binding (Table 3). However, we also uncovered examples of novel domain architectures that have not been previously described in other systems (Figure 5). One such architecture is found on six *Stentor* kinases (SteCoe_19985, SteCoe_20512, SteCoe_31546, SteCoe_35363, SteCoe_7134, and SteCoe_7398) and consists of a PAKA kinase domain with a FYVE domain near the N-terminus (Figure 5B). FYVE domains generally serve to target a protein into endosomal membranes, and have been observed on kinases of other families before [17], but this is the first example of a STE kinase with a FYVE domain. A more striking domain architecture, not previously described for any kinase, is found on two CDPKs (SteCoe_9200 and SteCoe_27913) with an adenylate kinase domain at the N-terminus (Figure 5A). Like other CDPKs, they also have EF hands at the C-terminus, and SteCoe_9200 possesses a fifth EF hand between the adenylate kinase and protein kinase domains.

**Figure 5.**
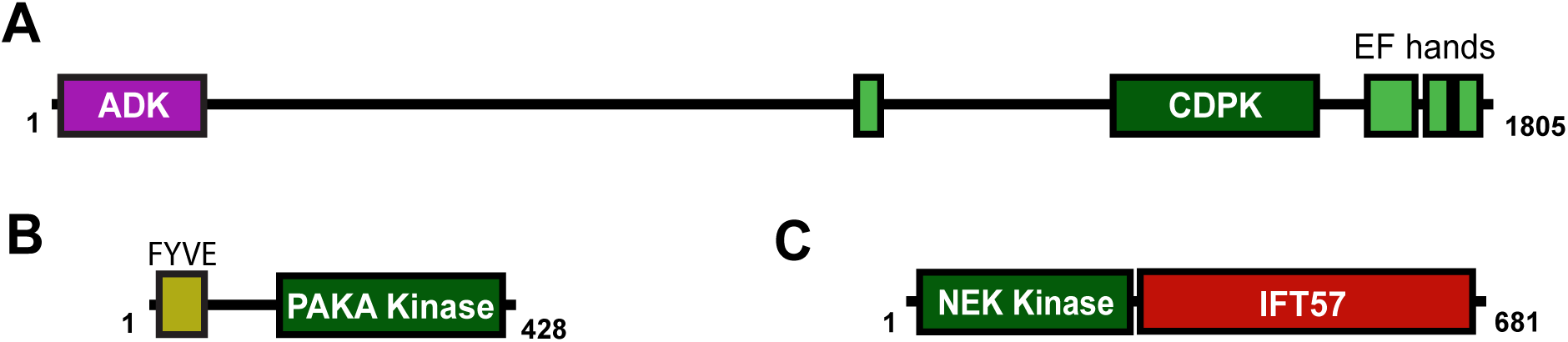
Novel domain architectures among Stentor kinases. **A:** Domain architecture of SteCoe_9200, a CDPK family kinase with an adenylate kinase domain at the N-terminus. Like most CDPKs, there are also EF hands at the C-terminus. **B:** Domain architecture of SteCoe_7134, a PAKA kinase with an N-terminal FYVE domain. **C:** Domain architecture of SteCoe_18659, an NEK family kinase with an IFT57 domain at the C-terminus. Numbers refer to amino acid positions. Drawn to scale.

**Table 3.** Other protein domains present on Stentor kinases.

| Domain | # | Domain | # | Domain | # | Domain | # |
| --- | --- | --- | --- | --- | --- | --- | --- |
| EF-hand | 287 | FYVE | 6 | ADK | 2 | DEP | 1 |
| cNMP binding | 97 | HEAT | 5 | Cyclin C | 2 | DUF547 | 1 |
| Ankyrin | 93 | LRR | 5 | Cyclin N | 2 | DUF3543 | 1 |
| POLO box | 43 | FAT | 4 | DUF3385 | 2 | NUC194 | 1 |
| Response regulator | 43 | WD40 | 4 | F-box | 2 | RCC1 | 1 |
| HATPase C | 42 | FATC | 3 | IFT57 | 2 | Ribonuclease 2-5A | 1 |
| TPR | 14 | Myosin_TH1 | 3 | Rapamycin binding | 2 | ubiquitin | 1 |
| zf-MYND | 9 | NAF | 3 | RWD | 2 | VWA | 1 |
| PH | 8 | PG binding | 3 | ANTH | 1 |  |  |

We also identified two NEK kinases (SteCoe_18659 and SteCoe_22983) with an IFT57 domain (Figure 5C). IFT57 is a protein involved in intraflagellar transport (IFT) [62]. It helps stabilize the IFT-B complex [63, 64], and the human homolog HIPPI is involved in the regulation of apoptosis, as well as cilia assembly and sonic hedgehog signaling [65, 66]. In most systems IFT57 domains are not found on kinases, though the domain architecture found in *Stentor* is also found in other ciliates. Perhaps the IFT57-like kinases of ciliates may provide more clues about the role of IFT57 in IFT and its interactions with other proteins.

Similarly, seven of the Aurora kinases in *Stentor* possess an N-terminal zf-MYND domain, which can mediate protein-protein interactions. While this domain architecture has not been observed on Aurora kinases from other eukaryotes, it is found in other ciliates.

### Pseudokinases

Active kinases belonging to the ePK superfamily generally contain twelve conserved subdomains [4], but most eukaryotic proteomes also encode pseudokinases. These are proteins with domains that are highly similar to kinase domains but with one or more critical residue in the catalytic site altered [67]. This usually results in a lack of kinase activity, although there are cases of predicted pseudokinases that have been experimentally shown to have some level of kinase activity [68–71]. However, even when a pseudokinase is known to be catalytically inactive, often the protein is still important to cell physiology, for instance in allosteric regulation of an active kinase or in scaffolding of an enzyme complex [67, 72, 73]. Substitutions in the lysine and aspartate residues of the active site motifs VAIK, HRD, and DFG are often used to predict pseudokinases, and studies suggest that pseudokinases tend to make up about 10% of the kinome in humans, about 11% in *Dictyostelium*, and about 22% in *Tetrahymena* and *Paramecium* [5, 8, 16]. We searched *Stentor* kinases for changes in one of these three highly conserved residues. The atypical protein kinases were not included in this search, since their kinase domains are significantly different from the well-defined ePK protein kinase domain fold. Of the 1977 *Stentor* ePKs, 289 (14.6%) were found to have a substitution in one of these three residues. Twelve of these were homologs of Scyl or Wnk. Scyl is a family found across eukaryotes composed entirely of pseudokinases; Wnk kinases are generally active but lack the conserved lysine residue [67].

We further examined whether any individual kinase families had higher rates of active site substitutions in *Stentor* (Figure 6A). A large number of the pseudokinases were unique or in *Stentor*-specific families (39 and 47, respectively), indicating a high degree of divergence across other sites in these genes.

**Figure 6.**
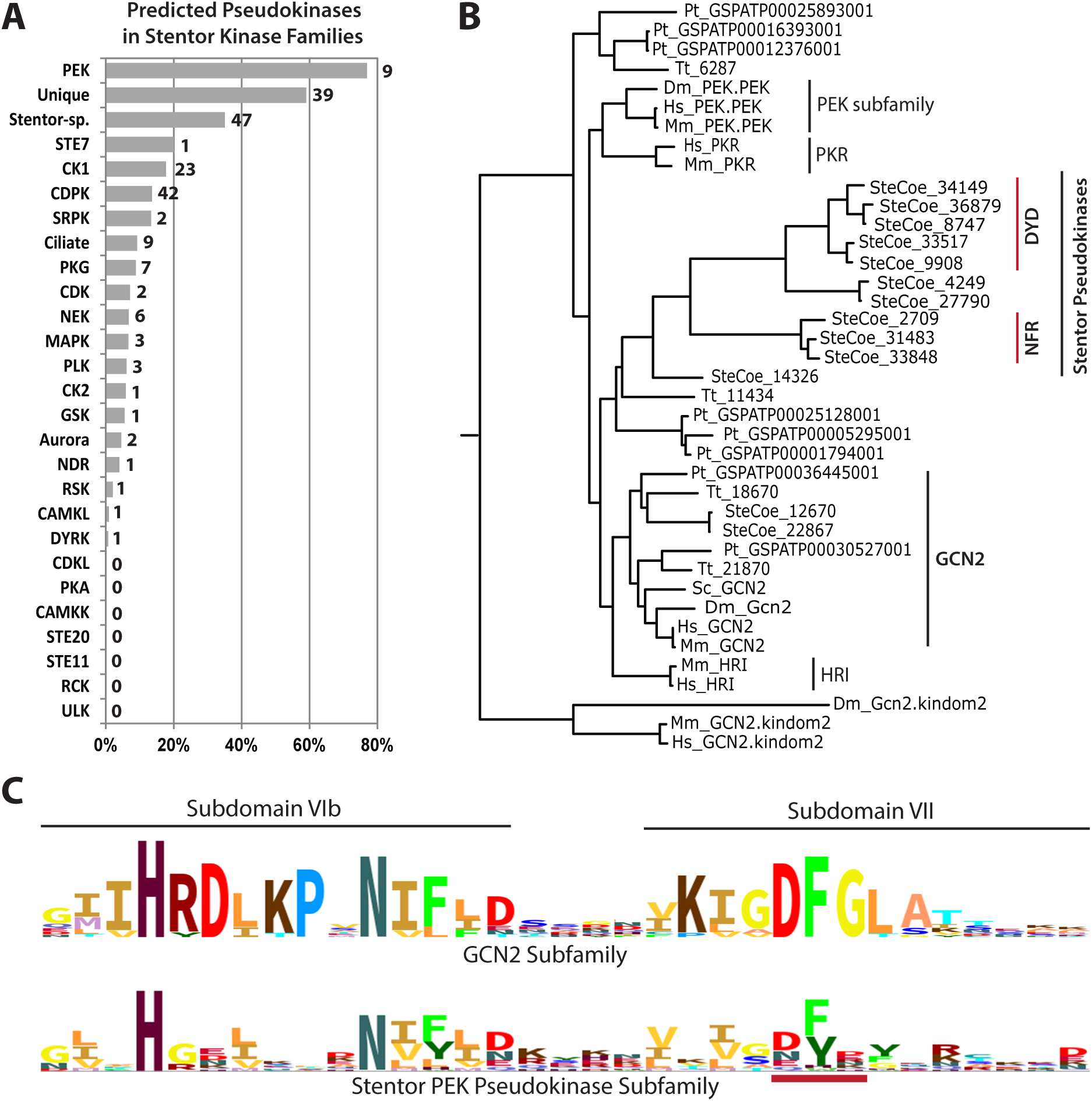
Pseudokinases among *Stentor* families. **A:** Proportions of pseudokinases are shown as a percentage of total family members in *Stentor*. Numbers next to each bar indicate absolute number of pseudokinases for that family. “Stentor-sp.” refers to *Stentor*-specific families, “Ciliate” refers to ciliate-specific families, and “Unique” refers to unique kinases in *Stentor*. **B:** Maximum likelihood tree of PEK family kinases in ciliates and metazoa. Red lines next to *Stentor* pseudokinases indicate unusual motifs that replace the usual DFG motif. Species represented as follows: SteCoe – *Stentor*; Tt – *T. thermophila*; Pt – *P. tetraurelia*; Sc – *S. cerevisiae*; Hs – *H. sapiens*; Mm – *M. musculus*; Dm – *D. melanogaster*. **C:** Comparison of HMM logos at kinase subdomains VIb and VII between the *Stentor* PEK pseudokinases and the similar GCN2 subfamily. Red line indicates position of motifs indicated on tree.

Of the kinases from conserved families, we found that the CDPK family in *Stentor* included 42 predicted pseudokinases out of 308 (13.6%) and the CK1 family included 23 out of 130 (17.7%). We note that the *Giardia* kinome contains 195 NEKs, a total of 70% of its kinome, and a surprising 71% of these were identified as pseudokinases [18]. Thus, there is a precedent for members of large kinase families evolving into pseudokinases, and perhaps these help regulate the active kinases.

Notably, we also found that the PEK family in *Stentor* contained 10 pseudokinases, representing 77% of the family. In the RNA-seq data from vegetative cells that helped inform the original gene models [3], these PEK pseudokinases have a mean of 1691 reads mapping compared to mean of 947 for the active PEKs (Additional File 1 Table S1), suggesting that they are expressed during vegetative growth and are not pseudogenes. Furthermore, maximum likelihood analysis suggests that they group together in a single subfamily (Figure 6B). In place of the normally-conserved DFG motif, we find most of these sequences either possess a DYD or NFR motif. Interestingly, sequence comparison between this subfamily and active PEKs reveals that while the HRDLKPxN and DFG motifs aren’t well conserved in this subfamily, the NIFLD motif in between is slightly more so (Figure 6C).

In contrast, other kinase families in *Stentor* exhibited much lower numbers of pseudokinases than the 14.6% average for the kinome. In particular, the DYRK family comprises an impressive 142 members in *Stentor*, but only one of these is a predicted pseudokinase. Additionally, several families contained no predicted pseudokinases at all (Figure 6A).

## Discussion

Here we have presented the kinome of *Stentor coeruleus*. We have identified 2057 protein kinases, and classified them into groups, families, and subfamilies. Many well-known kinases from other eukaryotes are conserved in *Stentor*, including MAPKs, PKAs and PKGs, as well as several mitotic kinase families including the NDRs, NEKs, PLKs, CDKs, and Auroras. We found many similarities to kinomes of ciliates and other protists, including the absence of PKC and the TK kinase group, and an expanded family of CDPKs.

In classical studies of *Stentor*, two properties tend to stand out: 1) its large size, and 2) its impressive regeneration ability. What signaling capabilities would be present in such a cell?

With respect to cell size, we note that generally the organisms that have kinomes of over 1000, mainly plants and ciliates, also have relatively large cells. Polyploidy in these organisms helps solve the need for larger scale protein production, but it is still easy to imagine that larger intracellular volumes require richer and more complex signaling repertoires. For example, *Stentor* has extensive and complicated cortical patterning. This is true of ciliates in general more than other phyla, but *Stentor*’s cortical patterning is more complex even than many other ciliates, especially within the oral apparatus. Oral regeneration alone requires the *de novo* assembly of tens of thousands of centrioles, with subsequent ciliogenesis. Accordingly, we find that some of the kinase families known to be involved in centriole assembly and ciliogenesis in metazoa, like PLKs and NEKs, are quite expanded in *Stentor*. It remains unclear whether the members in these families are functionally redundant, or whether they have evolved to exhibit greater specificity in localization or substrate.

Regarding regeneration, particularly of the oral apparatus, the process proceeds much like oral development during cell division. The cell division process even under normal conditions requires many steps, with formation of an entire new oral apparatus at the midpoint of the cell, where the anterior end of the posterior daughter needs to widen while the posterior end of the anterior daughter must constrict. The steps of cell division must be tightly regulated, and accordingly *Stentor* possesses many cell division kinases, including 28 CDKs and 44 Auroras.

Calcium signaling is also thought to play a role in regeneration [33]. In ciliates, calcium channels are present in the plasma membrane as well as intracellular structures like the contractile vacuole and the alveolar sacs, and calcium signaling has established roles in contraction and motility [29, 32, 74]. Furthermore, calcium is known to play a vital role in wound healing in metazoa [75–78]. It produces a clotting reaction in *Stentor* and has been hypothesized to play a central role in wound healing after cell cutting [79]. While other ciliates contain sizable CDPK families, the *Stentor* genome contains an impressive 308 CDPKs, making this the largest kinase family in *Stentor*, as well as the largest CDPK family described in an organism so far. Large CDPK families in ciliates in general may grant these large and complex cells greater versatility in calcium signaling capabilities. The need for this versatility may be even greater in *Stentor*, taking also into consideration its vast surface area and impressive wound healing capabilities. Two of the *Stentor* CDPKs even possess N-terminal adenylate kinase domains. This *Stentor*-specific innovation could serve to couple energy homeostasis with calcium signaling and highlights the potential for increased signaling versatility in large families.

Another kinase family known to be involved in regulating cell division and growth are the DYRK kinases. *Stentor* possesses 142 DYRK kinases, which is several fold higher than what is found in other ciliate genomes and appears to be the highest number of DYRKs encoded in a single genome to date. Furthermore, out of all of these, only a single one lacks one of the conserved residues previously shown to predict pseudokinases. This finding was striking, but previously studied DYRK kinases have been shown to self-regulate by autophosphorylation. Thus, it is likely there is selective pressure to retain a functional active site in order to maintain regulation.

*Stentor* has reductions in certain kinase families compared to other ciliates, though the reasons why are unclear. In particular, *Stentor* has fewer members of the ULK and HisK families than *Tetrahymena* or *Paramecium*, but even in these species the roles of these expanded families are not known. However, while these ciliates tend to have a lot in common with *Stentor* in terms of ultrastructure and lifestyle, there are important differences that set *Stentor* apart. In addition to *Stentor*’s much greater size, there are also behavioral disparities. For example, while *Tetrahymena* and Paramecium generally spend all their time swimming, *Stentor* possesses a sessile state and can easily transition between sessile and swimming behavior. As it spends less time swimming, perhaps *Stentor* has less need for vast quantities of kinases in families like the HisKs, which are very often involved in sensing extracellular environments.

## Conclusion

A kinome of over 2000 in *Stentor* underscores a rich signaling capacity which allows this single cell to maintain its large size, employ a variety of different behaviors, and even to regenerate after gross injury. It is characterized by multiple family expansions and contains both conserved and novel domain architectures, representing functional conservation as well as innovation within the kinases and distinguishing it even from other ciliates.

## Abbreviations

ePK: eukaryotic protein kinase
CaM: calmodulin
GC: guanylate cyclase
GPCR: G protein coupled receptor
HisK: histidine kinase
HMM: hidden markov model
IFT: intraflagellar transport
ML: maximum likelihood
TK: tyrosine kinase

## Additional Files

**Additional File 1. Table S1.** Classification of *Stentor* kinases. **Table S2.** Further classification of *P. tetraurelia* kinases. **Table S3.** *Stentor* non-kinase genes identified in this study: GPCRs, cyclins, and guanylyl cyclases.

**Additional File 2.** Nexus tree file of Stentor ePK Maximum Likelihood analysis.

## Acknowledgments

We thank Naomi Stover for hosting the Stentor genome database and for incorporating our gene model improvements. We thank Lydia Bright and Tatyana Makushok for critical reading of the manuscript, and for other members of the Marshall Lab for helpful discussions and comments.

## Availability of Data and Materials

All relevant data are published within the paper and its supporting additional files. Additionally, Stentor kinase gene annotations and updated gene models will be visible on StentorDB (stentor.ciliate.org).

## Authors’ Contributions

SBR and WFM conceived and designed the study. SBR analyzed data, and wrote the paper with help from WFM.

